# Inositol Polyphosphate-5-phosphatase K (*Inpp5k*) enhances sprouting of corticospinal tract axons after CNS trauma

**DOI:** 10.1101/2021.04.27.441184

**Authors:** Sierra D. Kauer, Kathren L. Fink, Elizabeth H.F. Li, Brian P. Evans, Noa Golan, William BJ Cafferty

## Abstract

Failure of CNS neurons to mount a significant intrinsic growth response after trauma results in chronic functional deficits after spinal cord injury. Approaches to identify novel axon growth activators include transcriptional and repressor screening of embryonic cortical and retinal ganglion neurons *in vitro*. These high throughput approaches have identified several candidates; however, their inability to comprehensively model the adult CNS has resulted in their exploitation *in vivo* failing to stimulate significant anatomical and functional gains. To identify novel cell autonomous axon growth activators while maintaining CNS complexity, we screened intact adult corticospinal neurons (CSNs) undergoing functional plasticity after unilateral pyramidotomy. RNA-seq of intact sprouting corticospinal tract (CST) axons showed an enrichment of genes in the 3-phosphoinositide degradation pathways, including six 5-phosphatases. We explored whether Inositol Polyphosphate-5-phosphatase K (*Inpp5k*) could enhance CST axon growth in clinical models of CNS trauma. Overexpression of *Inpp5k* in intact adult CSNs enhanced sprouting of intact CST terminals into the denervated cervical cord after pyramidotomy and cortical stroke lesion. *Inpp5k* overexpression also stimulated sprouting of CST axons in the cervical cord after acute and chronic severe thoracic spinal contusion. We show that *Inpp5k* stimulates axon growth by elevating the density of active cofilin in the cytosol of labile growth cones, thus stimulating actin polymerization and enhancing microtubule protrusion into distal filopodia. This study identifies *Inpp5k* as a novel CST growth activator and underscores the veracity of using *in vivo* transcriptional screening to identify the next generation of cell autonomous factors capable of repairing the damaged CNS.

**SIGNIFICANCE STATEMENT:** Neurological recovery is limited after spinal cord injury as CNS neurons are incapable of self-repair post trauma. *In vitro* screening strategies exploit the intrinsically high growth capacity of embryonic CNS neurons to identify novel axon growth activators. While promising candidates have been shown to stimulate axon growth *in vivo*, concomitant functional recovery remains incomplete. Using transcriptional profiling of intact adult corticospinal tract neurons undergoing functional plasticity, we identified *Inpp5k* as a novel axon growth activator capable of stimulating CST axon growth after pyramidotomy, stroke and acute and chronic contusion injuries. These data support using *in vivo* screening approaches to identify novel axon growth activators.

## INTRODUCTION

Spinal cord injury (SCI) results in chronic and debilitating deficits in sensory, motor, and autonomic function. CNS neurons fail to regenerate after injury due to a combination of their meager intrinsic growth potential (Curcio and Bradke, 2018) and the axon growth inhibitory environment of the adult CNS (Schwab and Strittmatter, 2014; Bradbury and Burnside, 2019). Overcoming both impediments will be critical to realizing significant functional recovery. Several potent anti-inhibitory strategies that target CNS myelin and chondroitin sulfate proteoglycan enriched extracellular matrix have been shown to elevate functional recovery after experimental SCI in rodents and non-human primates (Freund et al., 2007; Rosenzweig et al., 2019; Wang et al., 2020) and have entered clinical trials (Kucher et al., 2018). However, interventions that target the intrinsic growth potential of damaged and intact CNS neurons, while capable of stimulating robust axon growth, have been less successful in driving significant functional recovery after CNS trauma.

*In vitro* screening approaches to identify novel axon growth activators in CNS neurons have failed to translate into potent *in vivo* functional repair strategies (Moore et al., 2009; Blackmore et al., 2010; Buchser et al., 2010; Zou et al., 2015; Sekine et al., 2018). Disadvantages of *in vitro* screens include the embryonic age of the neurons and the simplicity of the *in vitro* growth environment, both of which poorly reflect the adult CNS. To overcome these challenges, two *in vivo* approaches have emerged to identify novel cell autonomous axon growth activators: identifying factors that drive the post-developmental growth of CNS neurons (Venkatesh et al., 2018; Venkatesh et al., 2021), and factors that support plasticity of intact CNS neurons after partial SCI (Fink et al., 2017). Recently we exploited the latter approach and profiled intact adult corticospinal tract neurons (CSNs) undergoing functional plasticity after unilateral corticospinal tract (CST) lesion (unilateral pyramidotomy, uPyX). Intersectional transcriptional profiling of mu-crystallin (crym)-GFP (Fink et al., 2015) expressing intact adult CSNs retrogradely labeled from the denervated spinal cord after uPyX revealed that plastic CSNs showed an enrichment in genes in the HIPPO, mTOR, 3-phosphoinositide degradation (3-PID) and LPAR1 (lysophosphatidic acid receptor-1) signaling pathways. We validated the pro-axon growth capacity of candidate members from each of these pathways *in vitro* and confirmed the efficacy of targeting the LPAR1 pathway to drive functional plasticity *in vivo*. This was the first study to probe the plasticity transcriptome and thus revealed a set of novel adult pro-axon growth activators. Here we sought to explore whether our lead candidate in the 3-PID, Inositol Polyphosphate-5-phosphatase (*Inpp5k*), could stimulate functional axon growth in CSNs after clinically relevant CNS trauma including stroke and spinal contusion injuries.

*Inpp5k* is one of six 5-phosphatases enriched in intact CSNs undergoing functional plasticity after PyX including *Fig4, Inpp5e, Inpp5j, Ocrl*, and *Synj1*, highlighting a potentially important role for this family of enzymes. The 10 mammalian 5-phosphatases have myriad roles include regulating protein trafficking, phagocytosis, synaptic vesicle recycling, neuronal differentiation, and neuronal polarity (Ooms et al., 2009). Together they terminate phosphoinositide-3-kinase (PI3K) signaling by hydrolyzing the 5 position on the inositol ring of phosphoinositides converting PI(3,4,5)P3 to PI(3,4)P2. PI3K signaling has an established role in regulating actin cytoskeletal dynamics, implicating *Inpp5k* as a novel pro-axon growth activator (Zhang et al., 2018).

Here we show that overexpression of *Inpp5k* in intact CSNs significantly elevates growth of CST axons in the denervated side of the spinal cord after uPyX and unilateral cortical stroke lesion. Furthermore, overexpression of *Inpp5k* stimulates sprouting of CST axons in the cervical spinal cord after acute and chronic severe thoracic spinal contusion injury. *In vitro* analysis revealed that *Inpp5k* enhances axon growth by releasing plasma membrane bound cofilin, thus stimulating actin polymerization in labile axonal growth cones. These findings validate *in vivo* screening approaches to identify cell autonomous axon growth activators and sets the stage for additional spatial and temporal analyses to identify cell-type specific therapeutic interventions.

## MATERIAL AND METHODS

### Mice

C57Bl/6 male and female mice for tissue culture and *in vivo* experiments were ordered from Jackson Laboratories. To minimize the number of animals used while maintaining enough rigor to achieve our scientific objectives (Festing and Altman, 2002), we used freely available power analysis (Hedwig.mgh.harvard.edu/sample_size/js/js_parallel_quant.html) to estimate sample sizes. The experiments are powered at 90% based on the number of animals in each group, the SD as determined by our previous Pyramidotomy data for PyX and stroke experiments (Cafferty and Strittmatter, 2006) and BMS data for the contusion experiments (Cafferty et al., 2010), and a significance level of 0.05.

### Surgery

All procedures and postoperative care were performed in accordance with the guidelines of the Institutional Animal Use and Care Committee at Yale University.

### Cortical AAV infusion

Prior to surgery, AAV-*Inpp5k*-V5 + AAV-mCherry, and AAV-*Inpp5k*-V5 + AAV-Flex-GFP were mixed, such that the final concentration of each virus was 2.55×10^13^ viral particles/ml, and aliquoted to blind surgeon to treatment. At P54-56, mice received unilateral infusion of AAV into motor cortex (M1). AAV transduction of CSNs was completed as previously described (Fink et al., 2017). Briefly, burr holes were made over sensorimotor cortex using a dental drill and 5 micro infusions of 150 nl of a 1:1 viral mixture were made to a depth of 0.7 mm (co-ordinates, +1 mm to -1 mm posterior to bregma and 0.5 -1.5 mm lateral to bregma) using a Hamilton syringe and Micro4 infusion device to deliver a total volume of 375 nl of each virus (750 nl total). All animals received post-surgical antibiotics (Ampicillin, 100mg/kg subcutaneously) and analgesia (Buprenorphine 0.05mg/kg subcutaneously) for 2 days post lesion. Two weeks after cortical infusion, mice underwent PyX, stroke, contusion, or sham lesions (see below). For the chronic contusion group mice received contusions 28 days prior to injections. Twenty-eight days post PyX and stroke, 52 days post-acute contusion and 91 days post chronic contusion, mice were perfused with 4% PFA, brain and spinal cord dissected and post-fixed overnight at 4°C and embedded in 10% gelatin for subsequent immunohistochemical analysis.

### Unilateral pyramidotomy (PyX)

Adult C57Bl/6 mice (n = 30) were anesthetized with ketamine (100mg/kg) and xylazine (15mg/kg) and placed in a supine position, an incision was made to the left of the trachea, and blunt dissection exposed the occipital bone at the base of the skull. The occipital bone was removed on the left side of the basilar artery with blunt Dumont #2 forceps to expose the medullary pyramids. The dura mater was pierced with a 30-gauge needle and resected. The left pyramid was transected unilaterally with fine Dumont #5 forceps to a depth of 0.25 mm (n = 18) or exposed and left intact for sham lesion (n = 12). No internal sutures were made, and skin was closed with monofilament suture. All mice received post-surgical antibiotics (Ampicillin, 100 mg/kg subcutaneously) and analgesia (Buprenorphine 0.05 mg/kg subcutaneously) for 2 days post lesion. One mouse from the PyX group died under anesthesia, all other mice recovered uneventfully.

### Unilateral stroke

Cortical stroke was induced photochemically by modifying procedures previously described (Labat-gest and Tomasi, 2013). Adult C57Bl/6 mice (n = 35) were anesthetized with ketamine (100mg/kg) and xylazine (15mg/kg), and 200 μL of 20 mg/mL of Rose Bengal dye (4,5,6,7-tetrachloro-2′,4′,5′,7′-tetraiodofluorescein sodium salt) in phosphate buffered saline (PBS) was injected intraperitoneally. Mice were then positioned in a stereotaxic frame and a midline incision was made in the scalp exposing the left sensorimotor cortex. The periosteum over the skull was removed and a cold light source with a 4.5 mm aperture was centered 1 mm anterior and 0.5 mm lateral bregma. Five minutes after Rose Bengal injection, mice were illuminated for 15 minutes through the intact skull to induce a cortical stroke lesion (n = 20), or placement of the light source without illumination (control, n = 15). The scalp was closed with monofilament suture. Mice received antibiotics and analgesia as above. Two mice died from the control group died under anesthesia, all other mice recovered uneventfully.

### Severe contusion

Adult C57Bl/6 mice (n = 52) were anesthetized with ketamine (100mg/kg) and xylazine (15mg/kg). A skin incision was made along the midline of the back and thoracic vertebrae exposed via blunt muscle dissection. Laminectomy of the 9^th^ thoracic vertebra (T9) was completed, and the vertebral column stabilized using Adson forceps to grasp the lateral edges of the rostral (T7) and caudal (T10) vertebral body at the exposed foramen. The contusion probe was positioned 2 mm above the spinal cord and a 75 kilodyne (kdyne) impact force was delivered to the exposed spinal cord using Infinite Horizon impactor (Precision Systems Instrumentation) causing a severe contusion injury. Animals in the sham condition received laminectomy without impact. Animals received an internal suture and external sutures prior to being returned to their home cage which was placed on a heating pad. All animals received post-surgical antibiotics (Ampicillin, 100mg/kg subcutaneously) and analgesia (Buprenorphine 0.05mg/kg subcutaneously) for 2 days post lesion. One mouse in the sham group and 2 mice in the contusion group did not recover from anesthesia, all other mice recovered uneventfully.

### Histology and Immunohistochemistry

Mice were euthanized with CO_2_ and were transcardially perfused with 0.9% NaCl (normal saline) 10% heprin followed by 4% PFA in PBS. Brains and spinal cords were dissected, post fixed in 4% PFA overnight at 4°C and subsequently embedded in 10% gelatin (Sigma Aldrich) dissolved in water for vibratome sectioning. Transverse sections (35 μm) of cervical spinal cord (C6 -C7) coronal sections of brain and brainstem, and horizontal sections of thoracic levels (contused mice) were processed for mCherry with tyramide signal amplification (Perkin Elmer, Waltham, MA). Immunofluorescence utilized primary antibodies directed against mCherry (1:500,000, Abcam, USA), V5 (1:500, Sigma), GFAP (1:2000, DAKO) and PKCgamma (1:250, Santa Cruz Biotechnology, USA) and detected with secondary antibodies Alexa Fluor-488, -568, -647 (1:500, Life Technologies, Grand Island, NY). An investigator blinded to treatment and lesion completed all immunohistochemical procedures.

### Cortical Neuron Cultures

Embryonic cortical neurons were cultured as described previously (Fink et al., 2017). Briefly, E17 mouse embryos were dissected from timed-pregnant C57Bl/6 mice and cortices were rapidly dissected in ice-cold Hibernate E medium (Brain Bits). Cortices were then digested for 30 minutes at 37°C using a digestion medium containing papain (25 U/ml, Worthington Biochemical), DNAse I (2000 U/ml, Roche), 2.5 mM EDTA, and 1.5 mM CaCl2 in a neuronal culture medium composed of Neurobasal A (Life Technologies) with B27 supplement (Life Technologies), 1% sodium pyruvate (Life Technologies), 1% glutamax (Life Technologies), and 1% penstrep (Life Technologies). After digestion, cortices were washed twice with the neuronal culture medium and triturated in 2 ml of medium. Triturated cells were passed through a 40 μm cell strainer (Corning) and counted.

### Acute dissociated cultures for outgrowth assay

Dissociated cortical neurons were electroporated with AAV plasmid to overexpress a candidate gene or reporter control (either PMAX GFP or *Inpp5k*). Dissociated cortical neurons were added to 100μl of Nucleofector solution (Lonza) and 3μg plasmid DNA. Cell mixture was transferred to a cuvette and electroporated in a Lonza Nucleofector 2b device using the device’s pre-loaded mouse hippocampal neuron electroporation parameters. Electroporated neurons were then transferred to pre-warmed neuronal media and plated at a density of 2,000 cells per well in an 8-well glass Lab-Tek slide (Thermo Scientific) pre-coated with poly-D-lysine (Corning) and laminin (Life Sciences). For neuronal outgrowth assay neurons were cultured 4 days at 37°C prior to fixation.

### Scrape assay

Cells were plated on 96-well plates pre-coated with poly-D-lysine (Corning) at a density of 50,000 cells per well in 200 µl neuronal culture medium as described previously (Zou et al., 2015; Fink et al., 2017). Immediately after plating, 1 µl of AAV was applied to each well at a titer of 1.0×10^10^ viral particles/ml. Each plate contained wells that are given control AAV-YFP or AAV-mCherry virus to serve as a viral overexpression control for wells given AAVs to overexpress candidate genes. Neurons were then incubated at 37°C/5% CO_2_. On DIV 7 and 14, 100 µl of medium was removed from each well and replaced with 100 µl fresh neuronal culture medium. Prior to medium exchange on DIV 14, 96-well cultures were scraped using a custom-fabricated 96-pin array as described previously (Huebner et al., 2011; Fink et al., 2017). Neurons were placed back at 37°C and allowed to regenerate for 72 hours before fixation with 4% PFA in 4% sucrose. Regenerating axons were then visualized by staining for βIII-tubulin (1:2000, Promega). Cell density was visualized using nuclear marker DAPI (0.2mg/ml, Sigma) to ensure that each well contained the same number of neurons after viral treatment. Images of the center of each well were taken using a 10X objective on an ImageXpress Micro XLS (Molecular Devices). The scrape zone was then analyzed for the total length of axons regenerating into the scrape zone in ImageJ by an investigator blind to treatment conditions. For each well, the total axon length of regenerating axons into the scrape zone was normalized to the average the total axon length of regenerating axon from control wells on that individual plate to create a regeneration index. Data presented is the average regeneration index of all wells (minimum 27 wells/condition) from cultures obtained from 3 separate litters.

### Immunocytochemistry

Cultures for axon growth assay and growth cone analysis were fixed with 4% PFA, 30% sucrose in PHEM buffer (300mM PIPES, 125 mM HEPES, 50 mM EGTAm 10 mM MgCl_2_) for 20 min. Fixed cells were quenched with 0.1 M Glycine for 15 min and permeabilized with 0.1% Triton X-100 in PBS for 5 min. cells were washed three times in PBS then blocked at RT for 1 hr in 2% Bovine Serum Albumin (BSA). After blocking cells were incubated with the primary antibodies at 4 °C overnight. Cells were then washed three times in PBS and incubated with secondary antibodies at RT for 2 hrs. Electroporated neurons were identified by visualizing the control GFP or by staining for the V5 (1:500, Sigma) tag on *Inpp5k*. Cells used for growth cone analyses were stained for βIII-tubulin (1:1000, Promega), Phalloidin 568 conjugate (1:500, Biotum), Phalloidin 405 conjugate (1:60, Biotum), Cofilin (1:250, Abcam), Phospho-Cofilin (Ser3) (1:250, Cell Signaling), and EB3 (1:250, Abcam) and detected with secondary antibodies Alexa Fluor-488, -555 (1:500, Abcam), -405, -647 (1:250, Abcam)

### Quantification

#### Cervical sections

Transverse cervical spinal cord sections were cut from all experimental animals in all experiments to assess mCherry+ CST axon distribution. Images were analyzed using ImageJ software (NIH). To determine the total number of CST axons labeled per animal, images of the dorsal column or medullary pyramid were taken at 20X on an epifluorescent microscope (LeicaMicrosystems) from five randomly selected sections from each animal. A 25 x 25 μm grid was then overlaid on the images and the number of mCherry+ axons, in three randomly selected boxes from the grid, was counted. The average number of axons from these boxes was then scaled up to determine the total number of axons labeled. For axon density measurements low-power images (10X) of the dorsal ipsilateral, dorsal contralateral, ventral ipsilateral, and ventral ipsilateral quadrants were taken at 10X to determine the area occupied by mCherry+ axons. Borders were drawn around the dorsal and ventral grey matter, thresholded, and skeletonized to determine density of the area. To determine the number of midline crossing axons sections were analyzed using Leica software zoom. A line was drawn from the base of the dorsal columns through the ventral medial fissure. Individual mCherry+ axons crossing the midline were counted. Experimenters were blind to all conditions. Statistical analysis was performed using Prism (GraphPad Software). Groups were analyzed using two-way ANOVA with Bonferroni post-hoc tests for multiple comparisons between surgery and treatment groups.

#### Thoracic sections

Horizontal sections were cut from all contusion mice to assess dorsal column retraction from lesion. Only sections where the fasciculated dorsal column was visible were included for analysis (3-5 sections). Lesion size and retraction from lesion were measured using the ruler feature on an epiflourecent microscope (LeicaMicrosystems) software. The lesion was determined by the area surrounded by GFAP immunofluorescence. Lesion size was determined by measuring the rostral border of the lesion to the caudal border of the lesion. CST retraction from lesion border was determined by measuring the distance between the rostral lesion border and the fasciculate dorsal column (determined by mCherry signal). Analysis was performed by an experimenter blind to conditions. Statistics were analyzed using an unpaired two-tailed Student’s t test to determine differences between treatment groups.

#### Gridwalking

Experimental mice in PyX and stroke experiments were assessed for skilled motor function using the grid walking task (Starkey et al., 2005). Mice were placed on an elevated 45 x 45 cm metal grid with 2.5 x 2.5 cm squares with dark cardboard walls creating a perimeter to make the environment more comfortable for the animals. Mice were videotaped via reflection from an angled mirror placed under the grid. Mice were recorded and allowed to explore the grid for 3 minutes. An experimenter blinded to treatment and lesion scored videos for the percentage of impaired steps out of the first 50 steps taken for each limb. Impaired steps included a foot slip where the limb fell completely between the rungs or an incorrectly placed step where either the ankle or the tips of the digits were placed on the rung instead of proper grasping of the rung. Animals were acclimated to the grid and then tested the day before AAV infusion, the day before PyX and stroke, and days 3, 7, 14, 21, and 28 postlesional processing. Data are presented as average missed step ± SEM for both the impaired forelimb and hindlimb. Data were analyzed via repeated-measures ANOVA with Bonferroni post-hoc test for multiple comparisons of between subjects (surgery condition and treatment) and within subjects (day post injury).

#### Basso Mouse Scale

Mice that underwent contusion and sham lesions were assessed using the Basso Mouse Scale (BMS) (Basso et al., 2006). Two investigators blinded to treatments completed BMS scoring. Acute contusion mice received BMS scores the day prior to contusion, and days 3, 7, 14, 21, 28, 35, 42, 49, and 56. Chronic contusion mice received BMS scores the day prior to contusion and on days 3, 7, 14, 21, 28, 35, 42, 49, 56, 63, 70, 77, 84, and 91. Data are presented as average BMS ± SEM. Data were analyzed via repeated-measures ANOVA with Bonferroni post-hoc test for multiple comparisons of between subjects (surgery condition and treatment) and within subjects (day post injury).

#### Cell Culture Experiments

All analyses were performed by an experimenter blinded to conditions. Slides used for axon length experiment were imaged at 20X using an epifluorescent microscope (LeicaMicrosystems). Five images from 3 different wells from each biological n (3) were taken of individual neurons expressing GFP or *Inpp5k*-V5. The length of the longest axon (from soma to tip of axon) was measured using the Simple Neurite Tracer (SNT) ImageJ plugin. Data were analyzed using an unpaired two-tailed Student’s t test. For experiments comparing looped vs. extending neurons 5 growth cones were counted in 5 wells per biological n (3). Growth cones were visualized using βIII-tubulin and Phalloidin and were categorized as looped or extending. Percentage of extending neurons was analyzed using an unpaired two-tailed Student’s t test. For cofilin analysis high power 63X images were taken using a Zeiss 880 with Airyscan processing. Five images per biological n (3) were taken of three channels (cofilin, phosphor-cofilin, and phalloidin). Separate channels were analyzed using Image J. Each channel was autothresholded, and a border was then drawn around the phalloidin and the selection was restored onto the cofilin and phosphor-cofilin channels. The density for each channel was then measured. Percentage of phosphor-cofilin (phosphor-cofilin/phalloidin) was subtracted from the percentage of cofilin (cofilin/phalloidin) to determine the amount of active cofilin. Data were analyzed using an unpaired two-tailed Student’s t test. For analysis of microtubules in growth cones high power 63X images were taken using a Zeiss 880 with Airyscan processing. Five images per biological n (3) were taken of three channels (βIII-tubulin, EB3, and phalloidin). Each channel was autothresholded, and a segmented line was traced over each filopodia from base to tip in the phalloidin channel. The selection was restored onto the βIII-tubulin and EB3 channels and a plot profile was acquired for each (intensity X length). Filopodia length was converted into percentage and the average intensity ratio of EB3/βIII-tubulin across filopodia for each growth cone was determined. Data were analyzed via repeated-measures ANOVA with Bonferroni correction for multiple comparisons between groups (EB3/βIII-tubulin intensity) and within groups (filopodia length).

### AAV production

Custom AAVs were produced as previously described (Fink et al., 2017). AAV-mCherry construct was provided by Dr. In-Jung Kim (Yale University). This construct expresses mCherry under the CAG promoter followed by a WPRE enhancer and SV40 polyadenylation signal. To generate AAV-CAG-mCherry, Dr. Kim modified two AAV vectors (Addgene plasmid #18917 and #38044). DNA of #18917 was cut with BamHI and Eco RI, vector backbone was kept and ligated with mCherry insert. mCherry sequence was amplified by PCR using the DNA of #38044 as a template with BamHI and EcoRI at the end of the amplified product for ligation.

To develop the *Inpp5k* overexpression construct, mCherry was replaced with the coding sequence of *Inpp5k* (*Inpp5k* gene ID: 19062). *Inpp5k* was PCR amplified using Platinum Pfx DNA polymerase (Thermo Scientific) from P1 brain cDNA library obtained from Dr. In-Jung Kim (forward primer: CGACCGGTCCACCATGCAGCACGGAGACAGGAA, reverse primer: GCACCGGTGATCTGTGGCTCAGGCTCAT), we added a Kozak sequence and an Age1 restriction site to the forward primer, and an Age1 site to the reverse primer with stop codon deleted. Insertion of *Inpp5k* was completed with several steps: removal of mCherry, insertion of V5, and insertion of *Inpp5k*. mCherry was cut out of the AAV plasmid using BamHI and EcoRI, vector backbone, purified, and blunt cut (Qiagen quick blunting kit) to generate a blunted AAV vector. V5 was PCR amplified from a plasmid provided by Dr. Sourav Ghosh (Yale University) (forward primer: CGGATATCACCGGTGGTAAGCCTATCCCTAAC, reverse primer: CGGATATCTCACGTAGAATCGAGACCGAG) with the forward primer containing an EcoRV and AgeI site and the reverse primer containing an EcoRV site and stop codon for V5. V5 was cut with EcoRV and ligated to blunted AAV vector. AAV-V5 and PCR amplified candidate genes were then cut with AgeI and ligated together to generate AAV-CAG-*Inpp5k*-V5. AAV-CAG-FLEX-EGFP-WPRE (addgene:51502) served as control for all *in vivo* experiments.

### AAV synthesis

For AAV production, a triple-transduction method was used as has been described previously (Park et al., 2015). Briefly, HEK 293 cells were cultured in ten 15 cm plates and polyethylenimine (Polysciences) transfected with an AAV plasmid overexpressing a candidate gene, delta F6 helper plasmid (UPenn Vector Core), and a plasmid expressing AAV capsid 2/1 (UPenn Vector Core). Cells were harvested, pelleted, and resuspended in freezing buffer (0.15 M NaCl and 50 mM Tris, pH 8.0) 48-60 hours after transfection. For viral purification, cells were lysed by undergoing two freeze-thaw cycles followed by benzonase treatment (EMD Chemical) for 30 minutes at 37°C. Lysate supernatant was collected by spinning tubes in a swinging bucket rotor (Eppendorf) at 3700g for 20 minutes. Lysate supernatant was then added dropwise to the top of a centrifuge tube containing a 15%, 25%, 40% and 60% iodixanol step gradient. The gradient was spun in a Vti50 rotor (Beckman Coulter) at 50,000 rpm for 2 hours at 10°C. The 40% fraction was collected and added to an Amicon Ultracel 100K (Millipore) for buffer exchange to PBS. A small amount of virus was used to test viral titer using qPCR. Concentration of virus was repeated on smaller Amicon Ultracels (Millipore) until desired high titer was reached. Final purified virus was aliquoted and stored at -80°C.

## RESULTS

### Inositol Phosphate compounds are enriched in intact CST neurons undergoing injury-induced functional plasticity

In a recent study, we completed transcriptional profiling of intact CST neurons undergoing functional plasticity after contralateral pyramidotomy (PyX, **Fig. 1A**) in plasticity sensitized nogo recetor-1 knockout mice (*ngr1*^*-/-*^). Using laser capture microdissection 28 days after uPyX and 14 days after unilateral intraspinal injection of fast blue (FB) in *crym*-GFP transgenic mice, we completed bulk RNA sequencing of intact non-sprouting CST neurons (GFP+/FB-) and intact sprouting CST neurons (GFP+/FB+). Differential gene expression analysis revealed that 2738 genes were up regulated and 506 downregulated in intact sprouting vs. non-sprouting CST neurons (**Fig. 1A**). To gain biological insight into the functions of these significantly differentially expressed (SDE) genes, we used Ingenuity Pathway Analysis (IPA). As reported in our previous study (Fink et al., 2017), pathways consistent with actin cytoskeletal dynamics, axon growth, axon guidance, and synaptogenesis were found to be significantly dysregulated in CSNs undergoing functional plasticity (**Figs. 1B, C**). Additionally, we found that several pathways associated with Inositol Phosphate signaling were also dysregulated, specifically we found that six 5-phosphatases in the 3-phosphoinositide degradation (3-PID) pathway were among the most highly enriched genes in intact sprouting CSNs (**Fig. 1B**), including *Inpp5k*, Inpp5e, Inpp5j, Synj1, Ocrl, and Fig4. IPA also revealed that cytoskeletal functions were the most enriched Diseases and Biofunctions terms within our SDE gene set (**Fig. 1C**). These data suggest that inositol signaling is converging with the cytoskeleton in intact CSNs undergoing functional plasticity after contralateral PyX and thus form part of the molecular machinery driving axon growth in these neurons. We selected *Inpp5k* to probe the possible impact of the 3-PID pathway to stimulate axon growth *in vitro* and *in vivo*. To explore, we transfected embryonic 17.5 (E17.5) cortical neurons with vectors encoding either GFP or V5-tagged *Inpp5k*. After 4 days *in vitro* (DIV) neurons were fixed and stained with antibodies against GFP, Beta(III)tubulin and V5. Neurons transfected with *Inpp5k* grew significantly longer neurites compared to GFP controls (**Figs. 1D-F**). These data are consistent with our previous finding showing that *Inpp5k* enhances axon growth of acutely dissociated cortical neurons at 3 DIV (Fink et al., 2017). To explore whether *Inpp5k* was enhancing neurite outgrowth via mTOR, we completed the *in vitro* scrape assay (Zou et al., 2015; Fink et al., 2017; Sekine et al., 2018), including the mTORC1 inhibitor rapamycin in culture media after the *in vitro* injury. Neurons transfected with *Inpp5k* showed significantly more neurite growth into the scrapped area compared to mCherry transfected controls (Figs. 1G-K). Inclusion of DMSO (rapamycin diluent) +/-300 nM Rapamycin did not impact neurite growth of cells transfected with either mCherry or *Inpp5k*. These data show that *Inpp5k* is stimulating enhanced axon growth *in vitro* via an mTOR independent mechanism.

**Figure 1:**
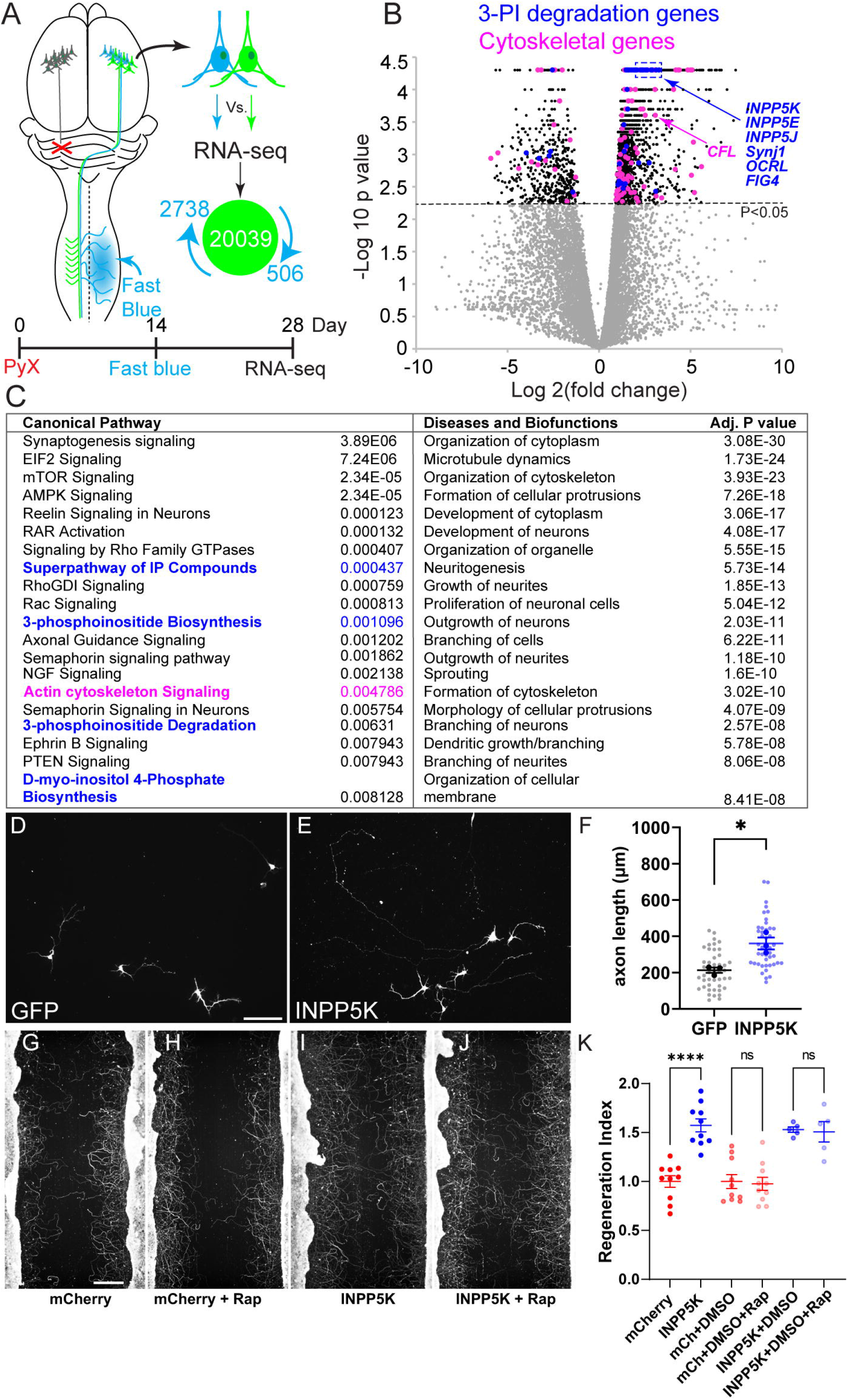
*Inpp5k* enhances neurite outgrowth and regeneration *in vitro*. Schematic (**A**) shows approach to identify intact CSNs undergoing functional plasticity after uPyX. Adult *ngr1*^*+/+*^ *crym*-GFP *ngr1*^*-/-*^ *crym*-GFP transgenic mice received a uPyX, 14 days post lesion mice received a contralateral infusion of the retrograde tracer fast blue, 28 days post lesion mice were prepared for laser capture microdissection of intact quiescent CSNs (GFP+FB-) and intact sprouting CSNs (GFP+/FB+) (Fink et al., 2017). Differential gene expression analysis calculated using a Wilcoxon rank sum test, showed that 2738 genes were significantly upregulated and 506 down regulated in sprouting vs. quiescent CSNs. Volcano plot (**B**)shows every gene profiled as log2 fold change versus –log10 of the false discovery rate (FDR) corrected p value. Non-SDE genes (grey dots) are separated from SDE genes (black dots) by *p* < 0.05 cut off (stippled) line. Inositol phosphate genes (dark blue) and genes associated with the cytoskeleton (magenta dots) are primarily enriched in sprouting CSNs. Five 5-phosphatases (*Inpp5k*, INPP5J, SYNJ1, OCRL, FIG4) and Cofilin are highlighted. Ingenuity Pathway Analysis (IPA) was conducted on genes enriched in CSNs undergoing functional plasticity. Table (**C**) shows the IPA output of significantly upregulated ‘Canonical Pathways’ and ‘Diseases and Biofunctions’ across the whole dataset of differentially expressed genes (with P values corrected for multiple comparisons), canonical pathways associated with inositol phosphate signaling are highlighted in blue and cytoskeletal dynamics in magenta. E17.5 neurons cultured transduced with *Inpp5k*-V5 (**E**), showed significant longer neurites compared to GFP (**D, F**, average total axon length of GFP (n = 45, grey dots) and *Inpp5k*-V5 (n = 45, light blue dots) neurons from n=3 independent experiments (*t*_4_ = 4.066, \**p* = 0.015, unpaired two-tailed *t* test). Data shown are the mean length of axon (µm) (biological n, darker dots) ± SEM. To assess if *Inpp5k* mediated enhanced neurite growth was mTOR dependent we transduced E17.5 cortical neurons with AAV-mCherry or AAV-*Inpp5k* +/-300 nM Rapamycin in DMSO (**G-J**). Neurons expressing mCherry showed minimal regeneration into the scrapped zone, while *Inpp5k* expressing neurons showed significant increase in regeneration compared to control. There was no significant difference in the neurite regeneration upon addition of rapamycin to mCherry or *Inpp5k* treated neurons (one-way ANOVA, \*\*\*\**p* < 0.0001 [F(5, 44) = 18.24], using *post hoc* Tukey’s HSD test). Scale bars, D = 100 µm, H = 200 µm.

### *Inpp5k* increases active cofilin and microtubule advancement in growth cones *in vitro*

Genes significantly differentially expressed in intact CSNs undergoing functional plasticity were enriched in pathways consistent with cytoskeletal rearrangements (Table **1C**). To explore whether *Inpp5k* was re-sculpting the axonal cytoskeleton and enhancing neurite growth we assessed the morphology of E17.5 cortical neuron growth cones (GCs) transfected with *Inpp5k* or GFP at 4 DIV. Strikingly, *Inpp5k* transfected neurons showed a significantly higher number of growth cones with an elongating or extending morphology compared to controls (**Fig. 2A-E**). While some GFP-transfected neurons displayed elongating GCs, the majority were in a looped, or stalled growth morphology at DIV4 (**Fig. 2E**). Previous studies have shown that a loss of active cofilin (non-phosphorylated form) decreases radially oriented filopodia, slows the advancement of microtubules into the periphery of GCs and thus increases the microtubule looping trajectories observed in stationary growth cones (Flynn et al., 2012). To explore whether *Inpp5k* was enhancing neurite growth via a similar mechanism, we examined the number of end binding protein 3 (EB3) comets extending into peripheral filopodia in *Inpp5k* and GFP-treated cultures. Significantly more EB3+ comets colocalizing with beta (III) tubulin+ microtubules were observed along the entire length of peripheral filopodia in *Inpp5k*-treated cultures compared to controls (**Figs. 2F-N**), indicating that an increased advancement of microtubules was driving the increased number of elongating growth cones after *Inpp5k* treatment. Furthermore, the density of active cofilin was increased in GCs after *Inpp5k* treatment compared to control (**Figs. 2O-W**). These data are consistent with the finding that *Inpp5k* has specificity for hydrolysis of PI(4,5)P2 (Yonezawa et al., 1990; Yonezawa et al., 1991; Ijuin et al., 2000) and that hydrolysis of PI(4,5)P2 results in a release and activation of a membrane-bound pool of cofilin (van Rheenen et al., 2007; van Rheenen et al., 2009). Taken together, these data indicate that the mechanism of increased axon extension in neurons overexpressing *Inpp5k* is via an increased available pool of active cofilin.

**Figure 2:**
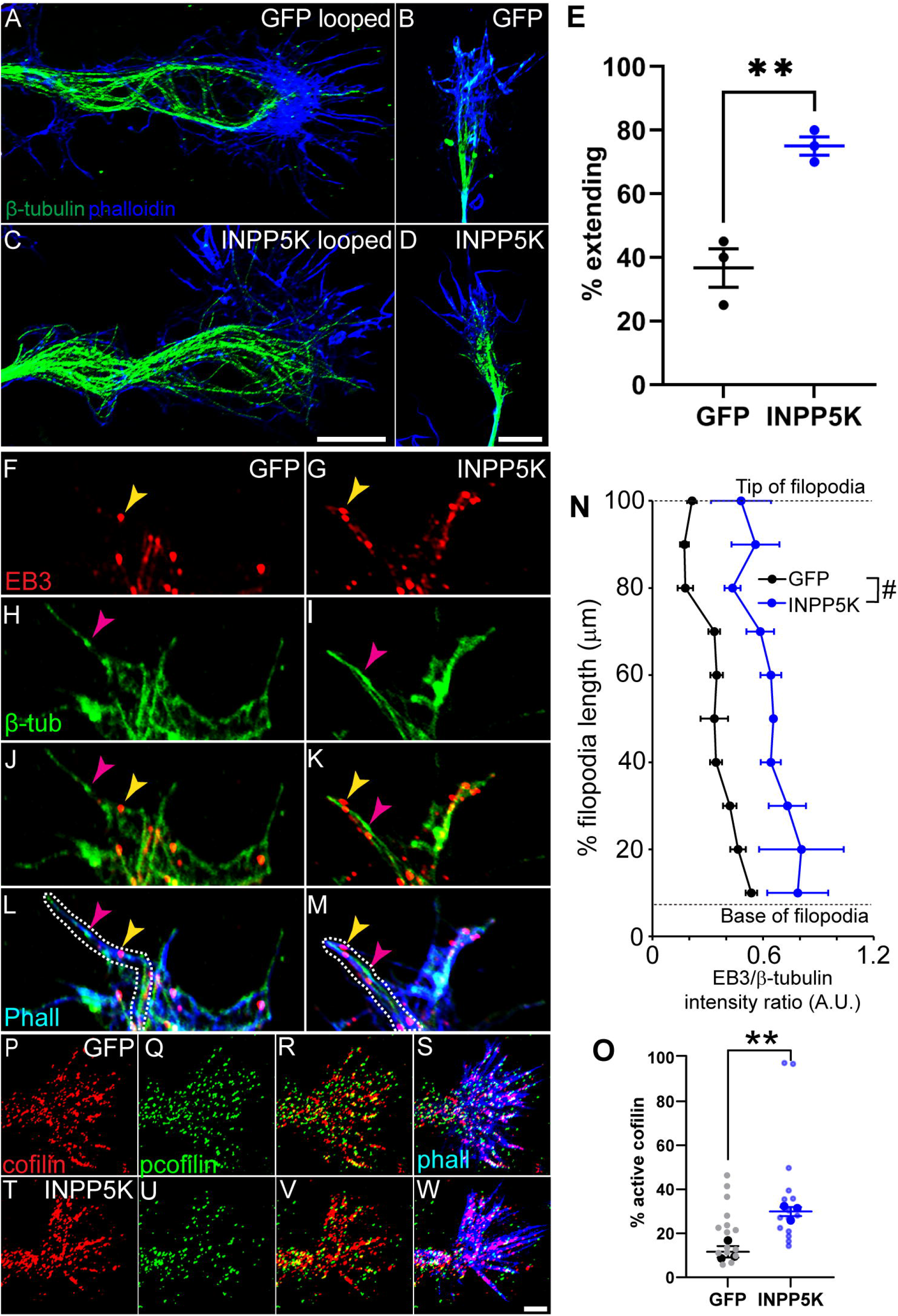
*Inpp5k* increases active cofilin and microtubule advancement in growth cones *in vitro*. Assessment of the morphology of growth cones (GCs) of E17.5 cortical neurons transduced with GFP (A, B) and *Inpp5k* (C, D) at 4DIV immunolabeled for β-tubulin (green) and phalloidin (blue), showed that significantly more GCs with extending or looped morphology was observed in *Inpp5k* treated cultures versus controls (E, GFP (n = 60) and *Inpp5k*-V5 (n = 60) neurons from n=3 independent experiments, unpaired two-tailed *t* test, *t*_4_ = 5.75, **p = 0.005, data are shown as the mean percent of extending growth cones ± SEM). Assessment of the relative filopodial location and density of EB3+ comments in GCs of E17.5 cortical neurons transduced with GFP (F, H, J, L, EB3 = red, beta-tubulin = green, Phalloidin = blue, pink arrows show co-localized EB3+ comets and microtubules, dotted line indicates the border of filopodia) and *Inpp5k* (G, I, K, M), shows that *Inpp5k* significantly elevated the density of EB3+ comets along filopodia (*F*(1, 4) = 10.21, *p* = 0.033, two-way ANOVA with repeated measures with Bonferroni *post hoc* comparisons), and the distance to which EB+ comets were found into distal regions of filopodia (*F*(1.339, 5.356) = 6.722. *p* = 0408, two-way ANOVA with repeated measures with Bonferroni *post hoc* comparisons). Data shown are EB3/β-tubulin intensity across percent of filopodia length ± SEM. Assessment of the relative density of (active) cofilin (red) and (inactive) phospho-cofilin (green) in GCs of E17.5 cortical neurons after 4DIV transduced with GFP (**P-S**) and *Inpp5k* (**T-W**), shows that *Inpp5k* treated cultures had significantly more active cofilin compared to controls (**O**, unpaired two-tailed *t* test, t(4) = 5.531, \*\**p* = 0.005). Data shown are mean density of active cofilin (biological n, darker dots, lighter dots = GCs, n=15/condition) ± SEM Scale bars, C = 20 µm; D = 10 µm; F = 4 µm; W = 4 µm.

### *Inpp5k* enhances plasticity of intact corticospinal tract neurons after unilateral pyramidotomy

As overexpression of *Inpp5k* increased axon growth of cortical neurons *in vitro*, we next sought to determine whether overexpression of *Inpp5k* could increase axon growth *in vivo* after CNS trauma. As our transcriptional screen (Fink et al., 2017) identified *Inpp5k* as enriched in intact CSNs after contralateral PyX, we initially sought to determine whether overexpression of *Inpp5k* in intact CSNs would enhance functional plasticity in this model. To explore, we co-infused either AAV-*Inpp5k*-V5 and AAV-mCherry or AAV-FLEX-GFP (control) and AAV-mCherry (**Fig. 3A**) into the right cortex of adult wild type (C57 Bl/6) mice two weeks prior to unilateral PyX (**Fig. 3A**). Twenty-eight days post-lesion, mice were euthanized and processed for histology. *Inpp5k*-V5 and mCherry are colocalized at the injection site (**Figs. 3B, C**) in layer 5 CSNs and in CST axons in the spinal dorsal columns (**Figs. 3D, E**), confirming that mCherry can be used as marker for transduced CSNs, their axons and their terminals.

**Figure 3:**
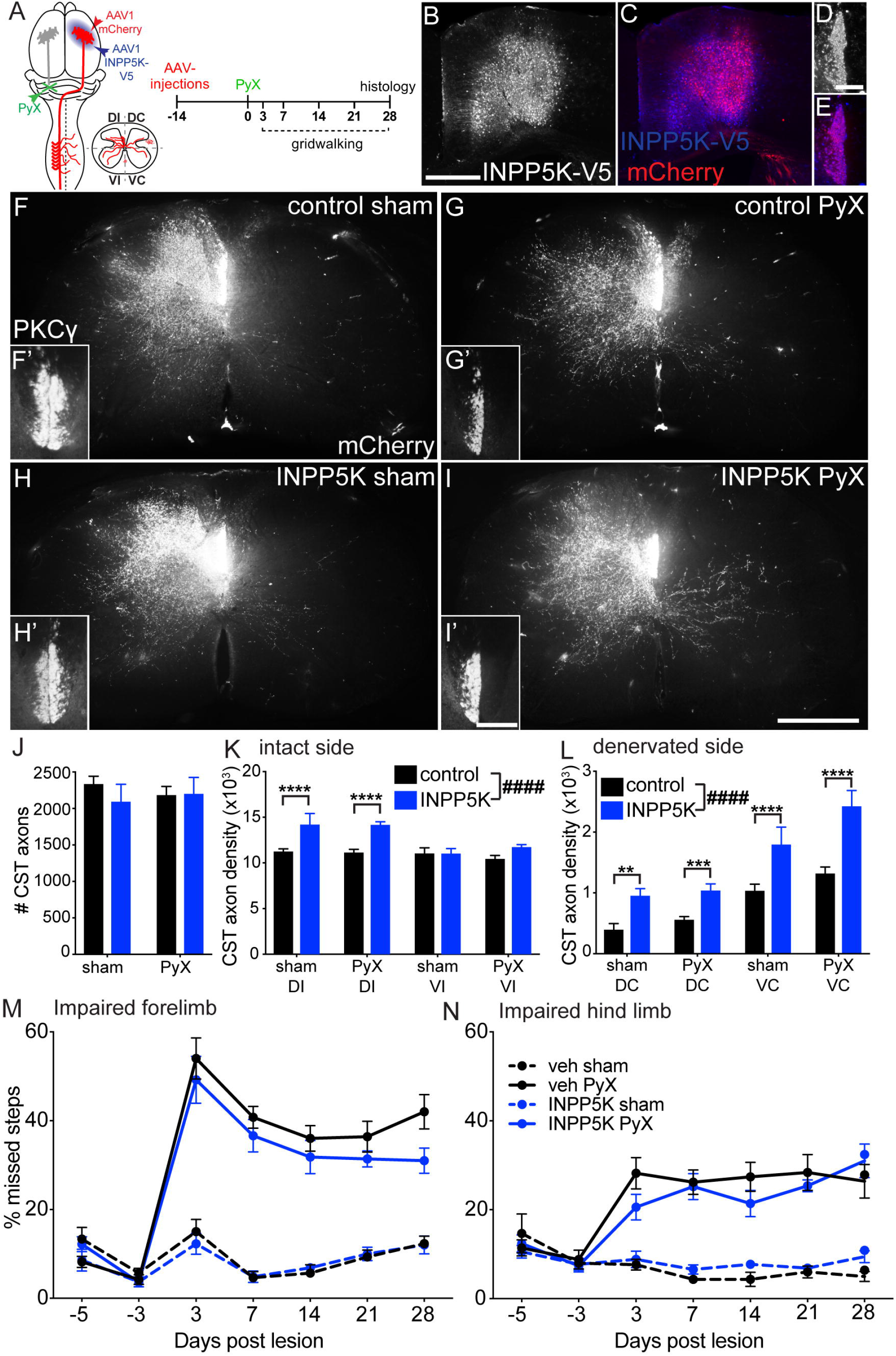
*Inpp5k* increases sprouting following PyX. Schematics (**A**) show relative locations of PyX and cortical injection of AAV1 mCherry and AAV1-*Inpp5k*-V5. Lower right schematic of transverse cervical sections with mCherry (red) labeling in four quadrants that were analyzed and right shows experimental timeline. Photomicrographs **B-E** show V5 (*B, D*) and mCherry (*C, E*) staining in M1 cortex (**B, C**) and C4 spinal cord 42 days after AAV infusion, confirming expression of *Inpp5k* and the reporter in CSNs and CST axons. Photomicrographs **F-I** show mCherry+ CST axon staining in transverse sections of C6 spinal cord from mice that received AAV1-FLEX-GFP after sham lesion (F) and PyX (G) and *Inpp5k* treatment after sham lesion (H) and PyX (I). Insets (**F’, H’**) show PKC_γ_ immunoreactivity both dorsal column projections in sham lesioned mice and intact contralateral dorsal columns after PyX (**G’, I’**). There was no significant difference in the number of CST axons traced between groups (**J**, two-way ANOVA with Bonferroni *post hoc* comparisons, *p* > 0.05, data are shown as average number of labeled CST axons ± SEM). Densitometric analysis of mCherry+ CST axons on the intact side of the spinal cord showed that there was a main effect of treatment, with *Inpp5k* treated mice having significantly more CST axon terminal density compared to controls (*F*(3, 50) = 18.97, *p* < 0.0001, two-way ANOVA with Bonferroni *post hoc* comparisons, n = 5 control sham, n = 11 control PyX, n = 6 *Inpp5k* sham, and n = 7 *Inpp5k* PyX). Specifically, *Inpp5k* treatment increased density of mCherry+ CST axons in the DI quadrant regardless of surgery condition (*F*(1, 50) = 34.69, *p* < 0.0001, two-way ANOVA with Bonferroni *post hoc* comparisons, data shown are average mCherry signal density ± SEM. (L) For the denervated side of the cervical cord *Inpp5k* treatment increased the density of mCherry+ CST axons regardless of surgical condition (*F*(3, 50) = 47.79, *p* < 0.0001, two-way ANOVA with Bonferroni *post hoc* comparisons), and this was true for both the DC and VC quadrants (*F*(1, 50) = 199.6, *p* < 0.0001, two-way ANOVA with Bonferroni *post hoc* comparisons, data shown are average mCherry+ signal density ± SEM. CST function was assessed using the grid-walking apparatus. No significant differences were found between *Inpp5k* PyX and control PyX treated subjects for the forelimbs (**M**, two-way ANOVA with repeated measures with Bonferroni *post hoc* comparisons, *p* > 0.05, data shown are average percent missteps ± SEM), or the hind limbs (**N**, two-way ANOVA with repeated measures with Bonferroni *post hoc* comparisons, *p* > 0.05, data shown are average percent missteps ± SEM. Scale bars, B = 1 mm; D = 100 µm; I = 500 µm; I’ = 100 µm.

CST axons expressing mCherry are present in the in the spinal dorsal and lateral columns, and throughout the grey matter contralateral to the cortical injection in transverse sections through C6/7 four weeks after uPyX (**Figs. 3F-I**). Counting mCherry+ puncta in the dorsal columns revealed no differences between lesion nor treatment among the groups (**Fig. 3J**) showing that *Inpp5k* did not affect the survival or health of CST axons. To analyze CST sprouting on both the intact and deneverated sides of the spinal cord, we partitioned the cord into four quadrants dorsal ipsilateral (DI), dorsal contralateral (DC), ventral ipsilateral (VI), and ventral contralateral (VC) (schematized in **Fig. 3A**). Laterality is described in reference to the lesion, therefore, as the CST is crossed, dorsal ipsilateral refers to the dorsal quadrant on the intact side. There was a significant impact of *Inpp5k* treatment on the density of mCherry+ CST axons in on both the intact (**Fig. 3K**) and denervated side (**Fig. 3L**). Post-hoc analysis revealed that intact sham lesioned mice treated with *Inpp5k* showed a significant increase in the density of intact mCherry+ CST axons in both dorsal quadrants and the contralateral ventral quadrant in comparison to sham lesioned control mice (**Figs. F-H, K-L**). Similarly, pyramidotomized mice treated with *Inpp5k* displayed a significant increase in mCherry+ CST axon density in both dorsal quadrants and the contralateral ventral quadrant in comparison to lesioned control mice (**Figs. 3G-I, K-L**). PKCg-immunoreactivity (PKCg-IR) confirmed completeness of PyX in lesioned groups (**Figs. 3G’, I’**). No difference was seen in the number of mCherry+ CST axons crossing the midline between groups.

To explore the functional impact of *Inpp5k* overexpression in intact CSNs after contralateral PyX, CST function was assessed using an elevated grid-walking task. Neither control nor *Inpp5k* treated sham lesioned mice showed a deficit in post training proficiency in the ability of their forelimbs or hind limbs to navigate the grid at any time during the testing period (**Figs. M-N**). However, both *Inpp5k* and control treated mice showed a significant increase in the number of missed steps 3 days after uPyX, in both the forelimb (**Fig. 3M**) and the hind limbs (**Fig. 3N**) on the affected side. The number of missed steps observed by the forelimbs recovered modestly over time, however, the hind limbs maintained a steady functional deficit throughout the testing period. There was no significant difference in recovery between *Inpp5k* and control treated mice. Thus, these data show that *Inpp5k* overexpression leads to sprouting of intact axonal arbors in the ipsilateral and contralateral spinal cord.

### *Inpp5k* increases plasticity following stroke

We next sought to determine whether *Inpp5k* could impact functional plasticity of intact CSNs after a more clinically relevant and extensive CNS injury. To this end, we completed unilateral photothrombotic stroke lesions in primary motor cortex (M1, **Fig. 4A**) in wild type mice. Two weeks prior to stroke lesion, wild type mice received unilateral (right side) cortical injections of either AAV-*Inpp5k*-V5 and AAV-mCherry or AAV-FLEX-GFP (control) and AAV-mCherry and were functionally assessed using the grid walking task for 4 weeks post lesion (**Fig. 4B**). Photothrombotic stroke resulted in a reproducible ischemic lesion in M1 while minimally damaging subcortical structures (**Fig. 4C**). Contralateral cortical infusion of tracers remained unilateral and were not taken up and retrogradely transported by intracortical CST connections which could have impacted behavioral outcomes (**Fig. 4D**). Histological assessment of mCherry+ CST axons 28 days post lesion showed a consistent number of axons were labeled in the medullary pyramids irrespective of lesion and or treatment (**Fig. 4E**). Transverse sections of cervical cord showed mCherry CST+ axons in one side of the dorsal columns and throughout grey matter on the intact side of the spinal cord (**Figs. 4F-I**). PKCg-IR confirmed that the stroke was unilateral and complete in lesioned groups (**Figs. 4G’, I’**). There was no significant difference in the density of mCherry+ CST axons in the intact side of the spinal cord between mice that received stroke and or *Inpp5k* treatment (**Figs. 4F-I, J**). However, *Inpp5k* treatment significantly increased the density of CST axons on the denervated side, specifically in the ventral contralateral quadrant after stroke (**Figs. 4I, K**). Furthermore, number of CST axons crossing the spinal midline was also increased after stroke in the *Inpp5k* treated group (**Fig. 4L**). Fine motor skill assessment using the grid-walking task showed that stroke significantly impaired both the forelimbs and hind limbs on the affected side compared to sham lesioned mice (**Figs. 4M, N**). There was no significant difference in the spontaneous functional recovery observed between control and *Inpp5k* treated groups (**Figs. 4M, N**).

**Figure 4:**
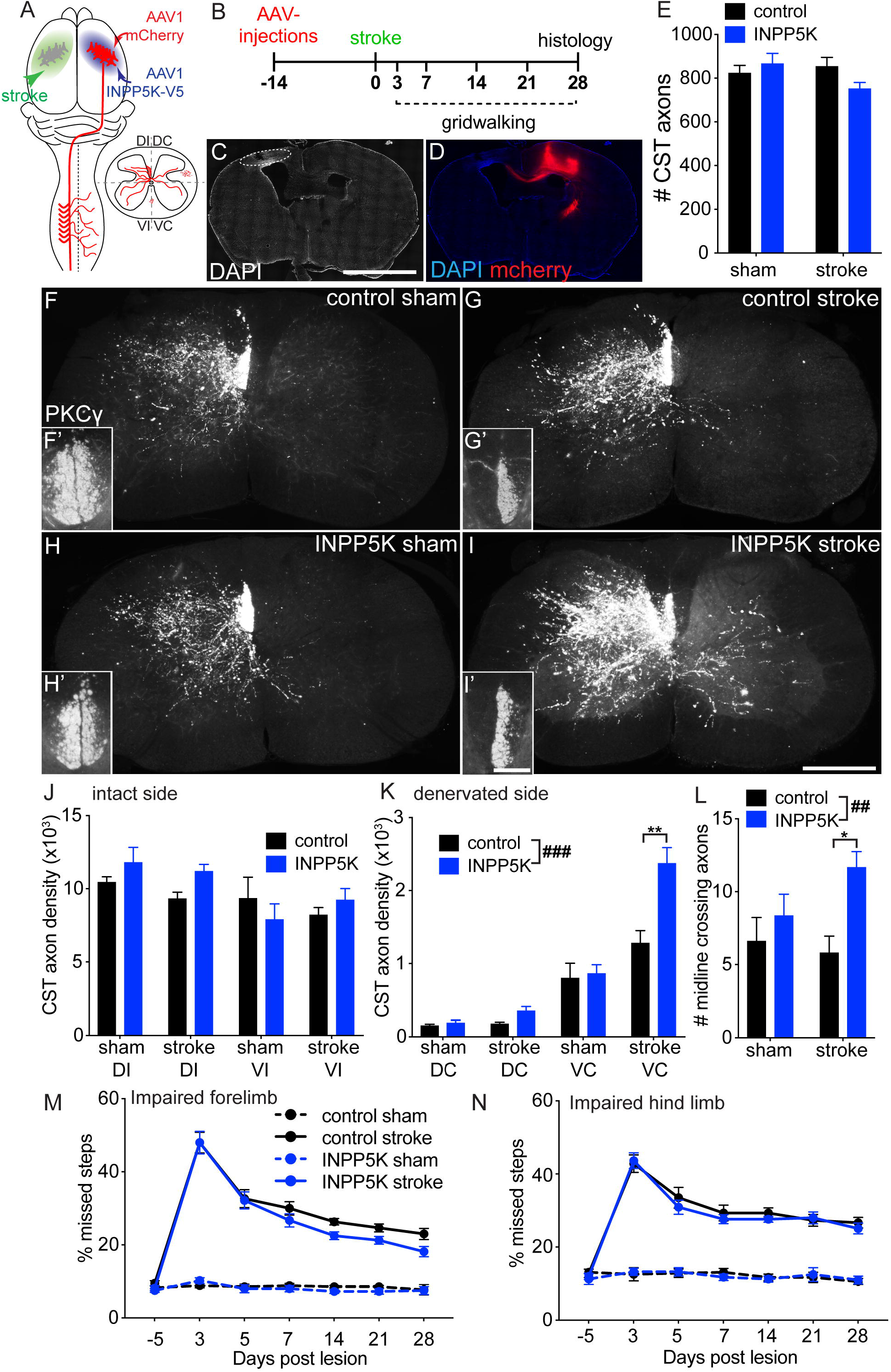
*Inpp5k* overexpression enhances sprouting of intact CSNs following stroke. Schematic (**A**) shows unilateral cortical stroke (green) and delivery of AAV1 mCherry and *Inpp5k*-V5 into contralateral cortex. Inset schematic shows quadrants of cord analyzed as before. Experimental timeline is shown in (**B**). Photomicrographs show accumulation of DAPI+ nuclei delineating the stroke the location (**C**, stippled oval) in the motor cortex, and mCherry+ CSNs in contralateral cortex 4 weeks after lesion (**D**). The number of mCherry+ CST axons counted in the C4 was invariant between group (**E**). Photomicrographs F-I show transverse sections through C6 showing mCherry+ CST axon labeling in control sham (**F**, n = 6), control stroke (**G**, n = 11), *Inpp5k* sham (**H**, n = 7), and *Inpp5k* stroke (**I**, n = 9) 4 weeks post lesion. Insets (F’-I’) show PKC_γ_-IR revealing complete unilateral denervation of CST in (G’, I’) stroke subjects and intact dorsal columns in (F’, H’) sham subjects. Densitometric analysis revealed that there was no significant effect of surgery or treatment on mCherry+ CST sprouting on the intact (ipsilateral) side (**J**, two-way ANOVA with Bonferroni *post hoc* comparisons, *p* > 0.05, data are shown as average mCherry + signal density ± SEM). *Inpp5k* treatment significantly enhanced CST sprouting on the denervated (contralateral) side compared to control (**K**, *F*(3, 58) = 7.817, ###*p* < 0.0001, two-way ANOVA with Bonferroni *post hoc* comparisons). Specifically, *Inpp5k* treatment increased mCherry + signal density in the VC quadrant for *Inpp5k* treated stroke subjects (*F*(3, 58) = 4.849, *p* = 0.005, two-way ANOVA with Bonferroni *post hoc* comparisons). Data shown are average mCherry + signal density ± SEM. The number of mid-line crossing CST axons was also significantly elevated in mice treated with *Inpp5k* after stroke (**L**, *F*(1, 29) = 9.396, ##*p* = 0.005, two-way ANOVA with Bonferroni *post hoc* comparisons). Assessment of CST function using grid walking analysis revealed that there was significant differences between *Inpp5k* stroke and control stroke treated subjects in the percentage of missed steps for forelimbs (**M**, two-way ANOVA with repeated measures with Bonferroni *post hoc* comparisons, *p* > 0.05), and hind limbs (**N**, two-way ANOVA with repeated measures with Bonferroni *post hoc* comparisons, *p* > 0.05, data shown are average percent missteps ± SEM. Scale bars, C = 2 mm; I = 500 µm; I’ = 100 µm.

### *Inpp5k* increases plasticity following acute SCI

Based on our observations that *Inpp5k* increases sprouting of intact CSNs in two models of plasticity, we wanted to determine if *Inpp5k* overexpression could enhance functional regeneration and sprouting in a clinical model of SCI, specifically in an acute and chronic severe thoracic contusion model. To explore, we injected either AAV-*Inpp5k*-V5 and AAV-mCherry or AAV-FLEX-GFP (control) and AAV-mCherry into the right motor cortex of wildtype mice two weeks before a 75 kdyne contusion at T9 or sham lesion (**Fig. 5A**). Hind limb motor function was assessed for 8 weeks using the Basso Mouse Scale (BMS). Horizontal sections through mid-thoracic cord showed mCherry+ CST axons regenerating up to the penumbra of the GFAP enriched lesion site in both control (**Fig. 5B**) and *Inpp5k* treated (**Fig. 5C**) mice. No CST axons were seen penetrating through the lesion and entering the spinal caudal to the lesion site in either group. Analysis of tissue sparing around the lesion confirmed that there was no difference between groups (*t*1.706 = 11, *p* = 0.116, unpaired two-tailed *t* test). Furthermore, there was no significant difference in the retraction of the mCherry+ CST axon regeneration front between the two groups. These data confirm that *Inpp5k* is not stimulating regeneration of cut CST axons.

**Figure 5:**
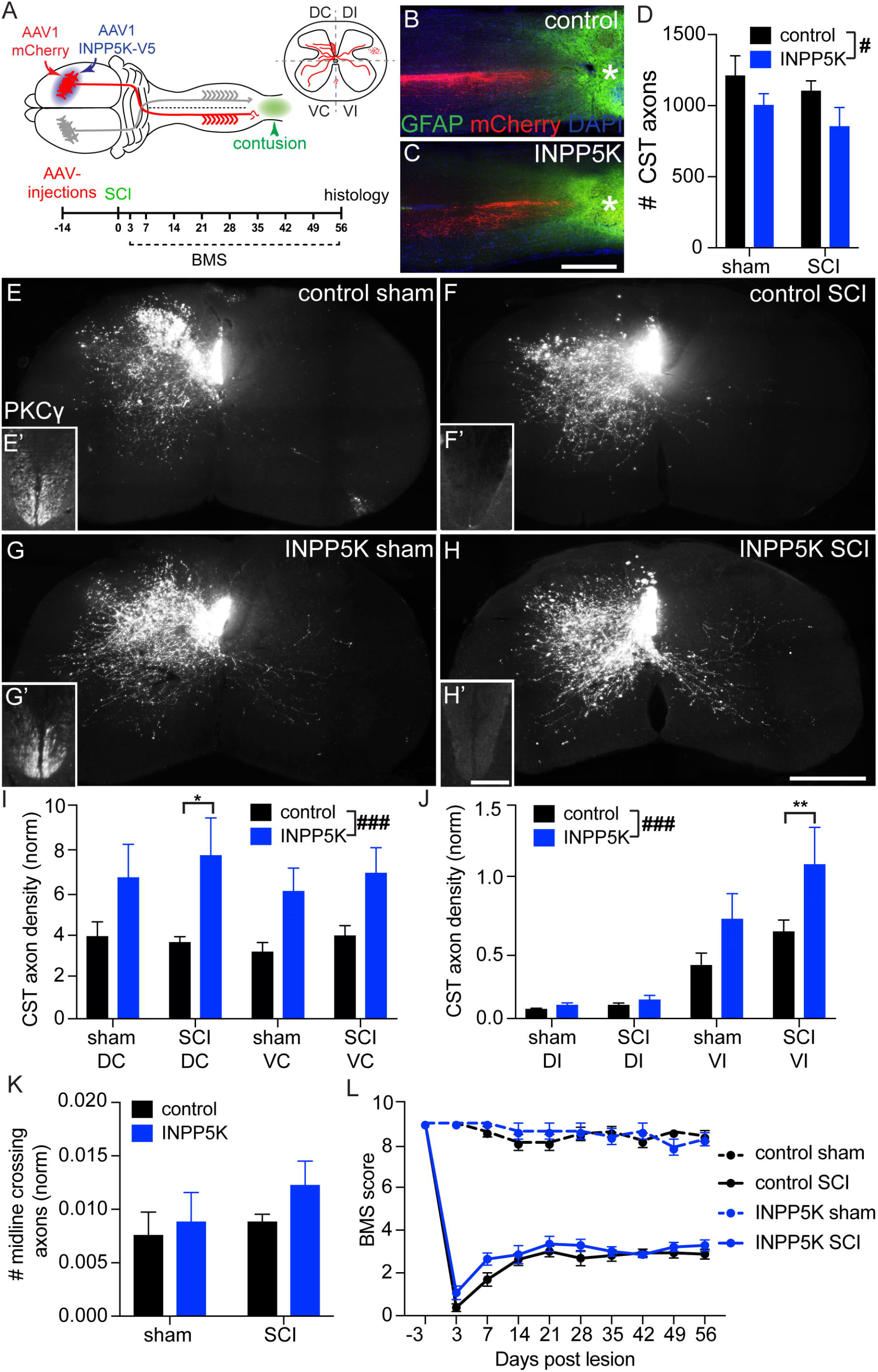
*Inpp5k* overexpression increases sprouting of CST axons after acute contusion injury. Schematic (**A**) shows bilateral T9 contusion injury (green) and delivery of AAV1 mCherry and *Inpp5k*-V5 into one side of motor cortex. Inset schematic shows quadrants of cord analyzed as before and experimental timeline. Photomicrographs show horizontal sections through the lesion site 4 weeks of severe contusion injury in control (**B**, mCherry+ CST axons in red, GFAP in green, asterisks indicate lesion site), and *Inpp5k* treated mice (**C**). No mCherry+ CST axons grew into or past the lesion site. The average number of mCherry+ CST axons counted in the dorsal columns was significantly different between groups (**D**, *F*(1, 21) = 5.215, *p* = 0.033, two-way ANOVA with Bonferroni *post hoc* comparisons, data shown are average number of mCherry + axons ± SEM), therefore subsequent densitometric data were normalized to # of CST axons per animal. Photomicrographs E-H show transverse sections through C6 showing mCherry+ CST axon labeling in control sham (**E**, n = 5), control SCI (**F**, n = 9), *Inpp5k* sham (**G**, n = 4), and *Inpp5k* SCI (**H**, n = 7). Insets show PKC_γ_-IR revealing intact lumbar dorsal columns for sham lesioned mice (**E’, G’**), and complete absence in lesion after SCI subjects (**F’, *H’***). Densitometric analysis of mCherry+ CST axons in grey matter showed that *Inpp5k* treatment significantly enhanced CST sprouting on the labeled (contralateral to cortical treatment, **I**, *F*(3, 42) = 7.753, *p* = 0.0003, two-way ANOVA with Bonferroni *post hoc* comparisons, data shown are average mCherry signal density normalized to number of dorsal column axons ± SEM), and on the non-labeled (ipsilateral to cortical infusion, **J**, *F*(3, 42) = 6.627, *p* = 0.001, two-way ANOVA with Bonferroni *post hoc* comparisons). There was an interaction of treatment and quadrant where *Inpp5k* SCI treated mice showed an increase in mCherry+ CST density compared to control in the ipsilateral quadrant control (*F*(3, 42) = 4.856, *p* < 0.006, two-way ANOVA with Bonferroni *post hoc* comparisons). There was no significant difference between surgery conditions and treatment groups for number of crossing axons (**K**, data shown are average number of mCherry axons normalized to number of dorsal column axons ± SEM). Behavioral assessment using the BMS revealed that there was no significance between *Inpp5k*, and control treated SCI mice (**L**, two-way ANOVA with repeated measures with Bonferroni *post hoc* comparisons, *p* > 0.05, data shown are average BMS score ± SEM). Scale bars, C = 2 mm; H = 500 µm; H’ =100 µm.

As we had observed significant plasticity of intact CST axons after PyX and stroke lesion, we sought to determine whether *Inpp5k* could stimulate axon sprouting in the cervical spinal cord after contusion injury. As we had treated and labeled only one side of cortex, we were able to complete similar analysis to our above PyX and stroke lesioned mice. Transverse sections of C6/7 spinal cord show mCherry+ CST axons in the left dorsal column and densely innervating one side of the spinal grey matter (**Figs. 5E-H**). PKCg-IR was observed bilaterally in the L4 segment of spinal cord in sham lesioned mice (**Figs. 5E’, G’**) and entirely absent from L4 in mice that underwent contusion (**Fig. 5F’, H’**). The number of CST axons labeled in the C4 dorsal columns was variable between groups and so densitometric data was normalized to the number of axons labeled per animal (**Fig. 5D**). *Inpp5k* increased mCherry density in the ipsilateral and contralateral quadrants in both sham and contusion lesioned mice (**Fig. 5I-J**).There was no difference in the number of mCherry+ CST axons crossing the midline (**Fig. 5K**). Furthermore, there was no significant difference in hind limb motor recovery between control and *Inpp5k* treated mice after contusion (**Fig. 5L**) using the BMS. Thus, *Inpp5k* overexpression enhances sprouting of intact or lesioned CST axons rostral to the severe contusion injury but did not stimulate regeneration of cut axons through the lesion site.

### *Inpp5k* increases plasticity following chronic SCI

To determine whether *Inpp5k* could stimulate sprouting and/or regeneration of CST axons after chronic contusion injury, we completed severe 75 kdyne T9 contusion injuries in wildtype mice, waited 28 days for hind limb motor recovery to plateau and then delivered either AAV-*Inpp5k*-V5 and AAV-mCherry or AAV-FLEX-GFP (control) and AAV-mCherry into the right motor cortex. Motor function was assessed with the BMS for an additional 9 weeks (**Fig. 6A**). Horizontal sections through mid-thoracic cord showed mCherry+ CST axons regenerating up to the penumbra of the GFAP enriched lesion site in both control (**Fig. 6B**) and *Inpp5k* treated (**Fig. 6C**) mice, however, no CST axons were seen penetrating through the lesion and entering the spinal caudal to the lesion site in either group. Analysis of tissue sparing around the lesion confirmed that there was no difference between groups (*t*1.037 = 13, *p* = 0.319, unpaired two-tailed *t* test). There was no significant difference in the number of CST labeled in the C4 dorsal columns between the groups (**Fig. 6D**) confirming that *Inpp5k* had no effect on the health or survival of chronically injured CST axons. To explore whether delivery of *Inpp5k* could stimulate sprouting of CST axons in the cervical cord 4 weeks after contusion injury, we completed identical densitometric analysis as above 13 weeks post lesion. Transverse sections of mCherry+ CST axons can be seen densely innervating one side of the spinal cord in both control (**Fig. 6E**) and *Inpp5k* treated mice (**Fig. 6F**). The absence of PKCg-IR in the L4 spinal cord confirmed that the contusion ablated the CST in its entirety (**Figs. E’, F’**). *Inpp5k* increased the density of mCherry+ CST axons in all quadrants on both sides of the cord compared to control treated mice (**Fig. 6G-H**). Additionally, *Inpp5k* treatment also increased the number of mCherry+ CST axons crossing the midline (**Fig. 6I**). However, hind limb motor scores using the BMS revealed that once mice had plateaued at 28 days, *Inpp5k* did not stimulate further motor recovery (**Fig. 6J**). In summary we found that *Inpp5k* overexpression enhances sprouting of chronically injured CST axons rostral to the lesion but did not stimulate regeneration of cut axons after chronic contusion injury.

**Figure 6:**
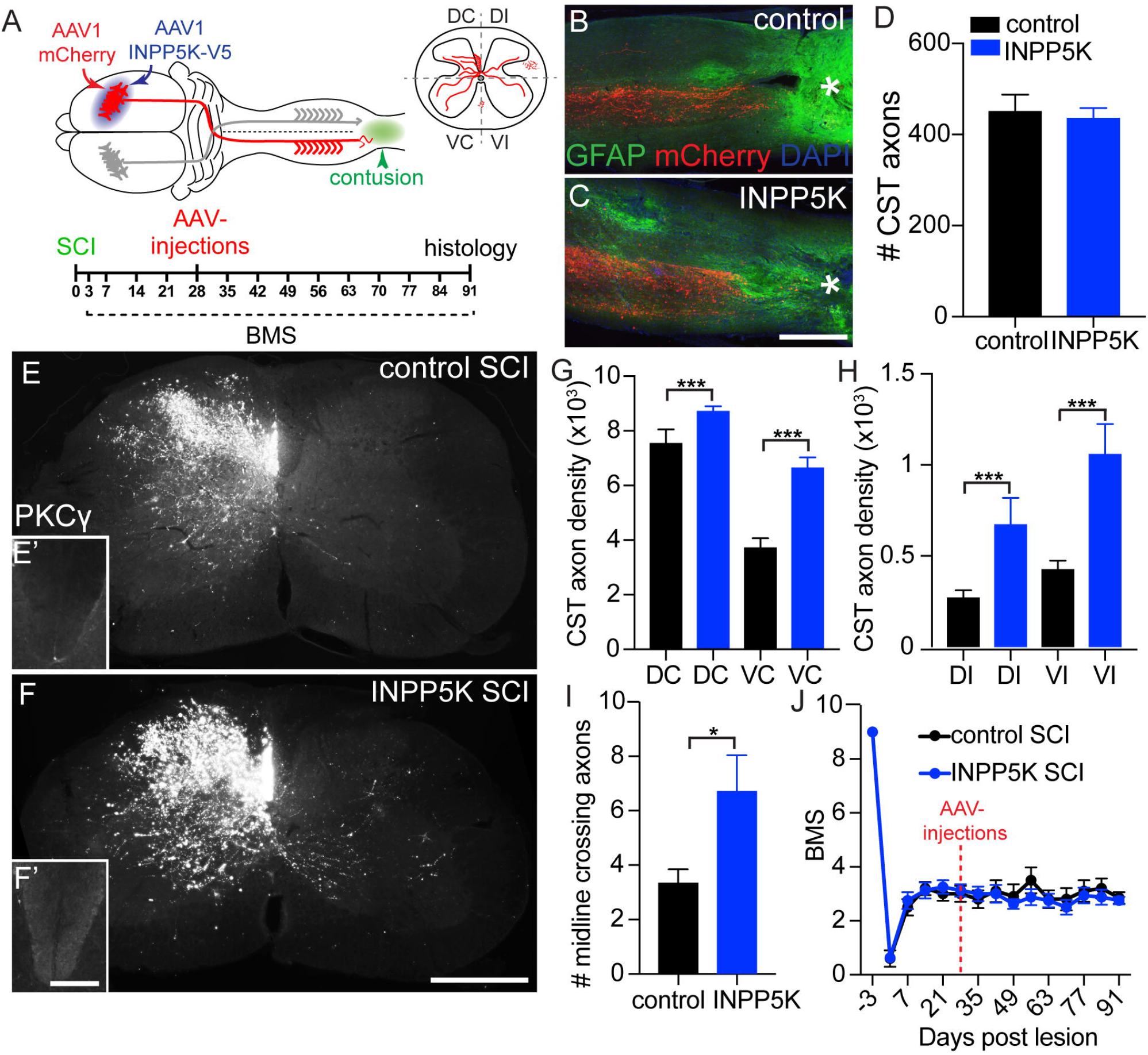
*Inpp5k* increases CST sprouting after chronic contusion injury. Schematic (**A**) shows bilateral T9 contusion injury (green) and delivery of AAV1 mCherry and *Inpp5k*-V5 into one side of motor cortex. Inset schematic shows quadrants of cord analyzed as before and experimental timeline, AVVs were delivered 28 days post SCI. Photomicrographs show horizontal sections through the lesion site 13 weeks of severe contusion injury in control (**B**, mCherry+ CST axons in red, GFAP in green, asterisks indicate lesion site), and *Inpp5k* treated mice (**C**). No mCherry+ CST axons grew into or past the lesion site. No significant differences were found in number of mCherry+ CST axons counted C4 the dorsal columns between treatment groups (**D**, data shown are average number of puncta ± SEM. Photomicrographs show transverse sections through C6 showing mCherry+ CST axon labeling in control SCI (**E**, n = 8), and *Inpp5k* SCI (F, n = 8). Insets show complete absence of PKC_γ_-IR in injured mice, **E’, F’**). Densitometric analysis of mCherry+ CST axons in grey matter showed that *Inpp5k* treatment significantly enhanced CST sprouting on the labeled (contralateral to cortical treatment, **G**, *F*(1, 29) = 25.33, \*\*\**p* < 0.0001, two-way ANOVA with Bonferroni *post hoc* comparisons, data shown average mCherry density ± SEM), and on the non-labeled (ipsilateral to cortical infusion, **H**, *F*(1, 29) = 19.53, \*\*\**p* < 0.0001, two-way ANOVA with Bonferroni *post hoc* comparisons). *Inpp5k* increased number of midline crossing CST axons compared to control (**I**, *t*14 = 2.348, *p* = 0.034, unpaired two-tailed *t* test, data shown are average number of axons ± SEM. Behavioral assessment using the BMS revealed that there was no significance between *Inpp5k*, and control treated SCI mice (**J**, two-way ANOVA with repeated measures with Bonferroni *post hoc* comparisons, *p* > 0.05, data shown are average BMS score ± SEM). Scale bars, C = 2 mm; F = 500 µm; F’ =100 µm.

## DISCUSSION

Transcriptional screening of intact adult CSNs undergoing functional plasticity revealed an enrichment of genes in the 3-PID. Here, we focused on the capacity of *Inpp5k* to stimulate CST axon growth after clinically relevant CNS trauma. In this study we show that: (1) *Inpp5k* increases axon outgrowth of cortical neurons *in vitro* by increasing the availability of active non-phosphorylated cofilin within labile axonal growth cones, (2) over-expression of *Inpp5k* enhances plasticity of intact CSNs after uPyX and unilateral cortical stroke, (3) *Inpp5k* increases sprouting of CSNs rostral to acute and chronic severe thoracic contusion SCI, (4) *Inpp5k* does not increase regeneration of damaged CSNs after SCI. From these data, we can draw several conclusions: first, *Inpp5k* is a novel pro-axon growth modulator in adult CSNs, second, confirmation that transcriptional profiling of adult CSNs undergoing functional plasticity is a robust approach to identifying novel cell autonomous axon growth activators, and, finally, that using combined localized retrograde tracing and transcriptional profiling may specifically identify pro-axon growth processes unique to a subset of neurons whose terminals are extending into the traced area, and are not generalizable axon growth activators, thus suggesting that comprehensive CST repair will require CSN sub-type specific interventions.

### *Inpp5k* mechanism of axon growth

INNP5K is part of the 3-phosphoinositide degradation pathway that is critical in modulating the activity of PI3K. Upon growth factor binding, PI3K catalyzes the conversion of PI(4,5)P2 to PI(3,4,5)P3 and via PDK1-dependent Akt phosphorylation, results in mTOR activation (Ooms et al., 2009). The PI3K/mTOR pathway controls cell survival and axogenic protein synthesis during development (Fonseca et al., 2014; Berry et al., 2016). Termination of PI(3,4,5)P3 synthesis is achieved via degradation to PI(4,5)P2 by the 3-phosphatase, *Pten. Pten* levels increase during development and into adulthood and restrict axon growth signaling via mTOR. Despite the potential for mTOR to cause deleterious effects (Berry et al., 2016), neuron-targeted deletion of *Pten* stimulates robust regeneration after SCI (Park et al., 2008; Liu et al., 2010; Ohtake et al., 2014). PI(3,4,5)P3 is also degraded by 5-phosphatases, including *Inpp5k* to produce PI(3,4)P2, which also activates PDK1/Akt (Ooms et al., 2009). Thus 5-phosphatases could elevate free PI(3,4)P2 and thus stimulate axon growth via mTOR. To explore this, we cultured embryonic cortical neurons overexpressing either control (mCherry) or *Inpp5k*, after 10 DIV culture wells were scrapped with a pin, and the media supplemented with the mTORC 1 inhibitor Rapamycin (**Fig. 1K**). Previously we had reported that rapamycin was capable of attenuating *Pten* inactivation-mediated enhanced neurite growth (Zou et al., 2015), thus establishing the veracity of this approach to determine a role for mTOR signaling in *Inpp5k* stimulating neurite growth. After 14 DIV, rapamycin failed to abrogate the *Inpp5k* effect. Thus, we concluded that *Inpp5k* was exerting its pro-growth effects independent of mTOR.

PI(4,5)P2 has also been shown to bind the actin polymerizing domain of cofilin, retaining it in the plasma membrane (Yonezawa et al., 1990; Yonezawa et al., 1991). For cofilin to participate in actin remodeling, it needs to be released from the plasma membrane. Indeed, release of cofilin from PI(4,5)P2 has been shown in a cancer cell line where EGF stimulated PLC released active cofilin into the cytosol where it bound severed F-actin and lead to actin polymerization and lamellipodia formation (van Rheenen et al., 2007). Crucially this study also showed that recombinant 5-phospatase activity was sufficient to release active cofilin into the cytosol. Active cofilin has previously been shown to drive neurite formation and elongation *in vitro* (Endo et al., 2003; Letourneau, 2009; Flynn et al., 2012; Dupraz et al., 2019) and shown to be necessary for regeneration of sensory neurons in the CNS *in vivo* (Tedeschi et al., 2019). Here we show that overexpression of *Inpp5k* in embryonic cortical neurons increased the density of active cofilin in growth cones (**Fig. 2O**) and enhanced neurite outgrowth. Thus, we propose that *Inpp5k* dephosphorylates PI(4,5)P2 releasing cofilin in its active state to promote actin remodeling (**Fig. 7A**) via either F-actin depolymerization or F-actin severing (**Fig. 7B**).

**Figure 7:**
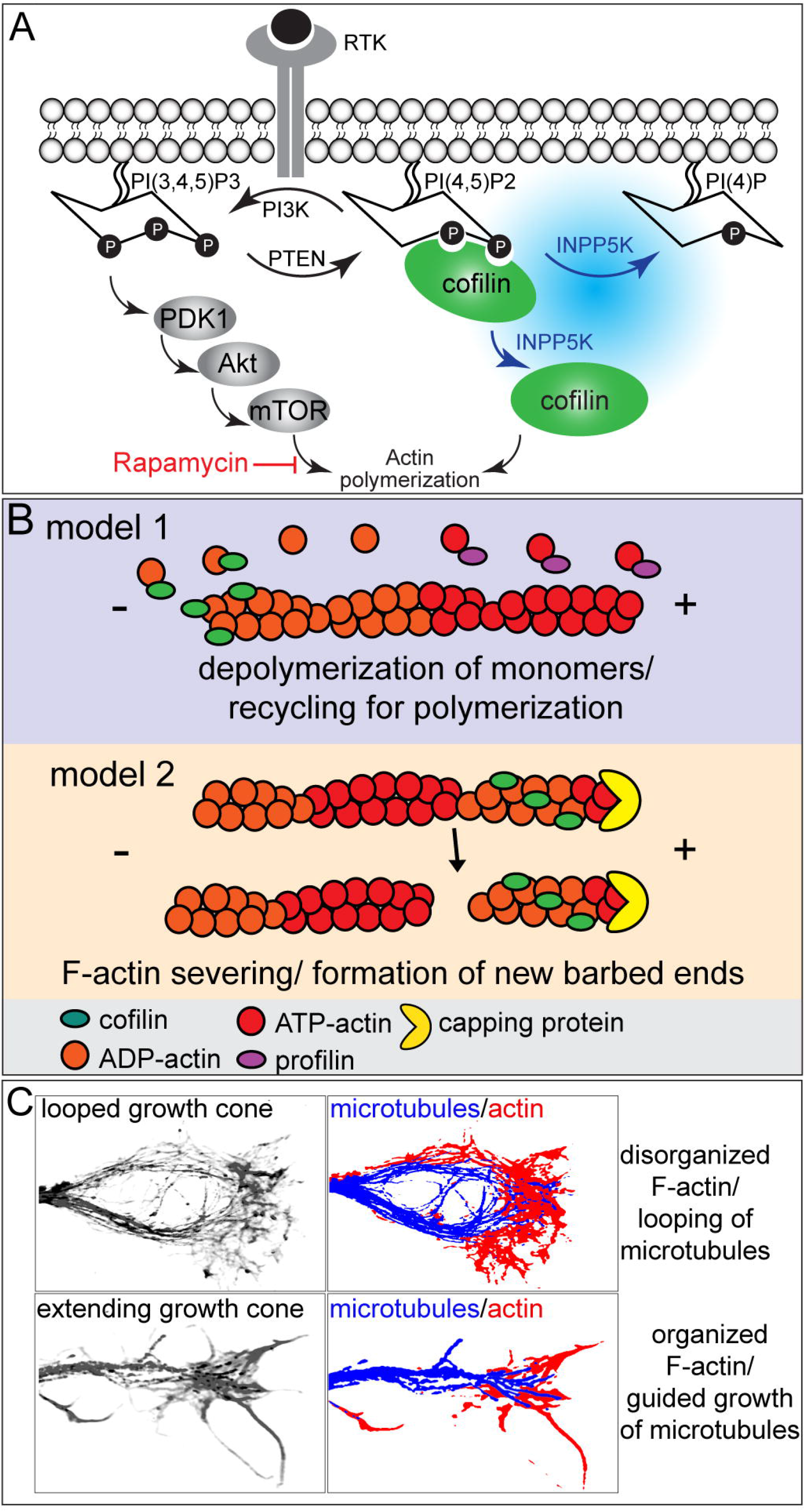
Model for mechanism of *Inpp5k* induced axon extension. PI3K is activated by growth factors binding to receptor tyrosine kinases phosphorylating PI(4,5)P2 forming PI(3,4,5)P3, initiating mTOR dependent (rapamycin sensitive) actin polymerization an enhanced neurite outgrowth. Dephosphorylation of basal levels of PI(4,5)P2 to PI(4)P by *Inpp5k* leads to unbinding of a pool of active cofilin from the plasma membrane that can engage in cytoskeletal remodeling (**A**) (Saarikangas et al., 2010). Two predominant hypothetical models by which cofilin regulates cytoskeletal remodeling leading to protrusion and extension of growth cones. In model one cofilin binds to ATP-actin at the pointed end of filaments, depolymerizing them. This increases availability of monomers converted to ATP-actin and polymerization by profilin. In model two capping proteins prevent polymerization. Cofilin severs F-actin creating new barbed ends for polymerization (**B**). LUT confocal images of paused looped growth cone (upper left) and extending growth cone (lower left). Right shows diagram of morphology of microtubules (blue) and F-actin (red). In the looped paused growth cone (upper right) F-actin is disorganized leading to abnormal microtubule growth. In the extending growth cone (lower right) F-actin is organized which directs microtubules in a radial orientation leading to directional growth.

The morphology of growth cones transfected with *Inpp5k* and GFP further supports this mechanism. *Inpp5k* overexpression prevented a looped morphology seen in paused growth cones in favor of extending growth cones, consistent with previous work (Flynn et al., 2012; Endo et al., 2003). F-actin microtubule interactions are an important driving force for microtubule orientation and extension of growth cones. Microtubules penetrate the growth cone periphery where they align with actin filament bundles (Dent and Kalil, 2001). Growth cones that are paused display a tightly bundled loop (**Fig. 7B, E**). Cofilin promotes severing of F-actin and the subsequent actin turnover allows the radial orientation of F-actin (Flynn et al., 2012). Microtubules associated with the actin filaments are then guided in an extending orientation. In the absence of cofilin, F-actin becomes disorganized, leading to looped and disorganized trajectories. *Inpp5k* expression increased polymerizing microtubules guided by filipodia in the periphery of growth cones. The microtubule protrusion and polymerization at the leading end of the growth cone leads to accelerated axon growth (Dupraz et al., 2019). Thus, we propose that *Inpp5k* increases axon extension via cytoskeletal remodeling and subsequently guided microtubule extension.

### *Inpp5k* enhances axon growth *in vivo*

Previously we showed that *Inpp5k* enhanced neurite outgrowth in DIV3 cortical neuron cultures (Fink et al., 2017). Here we explored whether *in vivo* overexpression of *Inpp5k* could elevate functional growth of intact and axotomized neurons in clinical models of CNS trauma. First, we assessed *Inpp5k* overexpression in intact CSNs after uPyX. *Inpp5k* stimulated significant sprouting of intact CST arbors throughout spinal grey matter on the intact and denervated side of the spinal cord. Despite robust sprouting into the denervated ventral horn, *Inpp5k* treated mice did show an increase in functional recovery versus controls. These observations are consistent with studies that target monogenic manipulation of intact CSNs including *Lppr1* overexpression (Fink et al., 2017), *Klf6* overexpression (Kramer et al., 2021), and *Pten* deletion (Geoffroy et al., 2015) that show increased sprouting of intact CSNs post uPyX, but not elevated functional recovery. We observed a similar result after cortical stroke, *Inpp5k* overexpression enhanced sprouting of intact CSNs into the denervated side of the spinal cord without enhancing functional recovery. Without additional stimuli that can guide labile arbors to correct postsynaptic targets, such as rehabilitative training (Wahl et al., 2014; Serradj et al., 2017; Loy et al., 2018; Torres-Espin et al., 2018; Loy and Bareyre, 2019), it is not surprising that significant functional recovery is not observed after monogenic intervention.

To assess whether *Inpp5k* could also stimulate regeneration of lesioned CST axons, we overexpressed *Inpp5k* or FLEX-GFP (control) in adult mice either 2 weeks (acute treatment) prior to, or 4 weeks post (chronic treatment) a severe bilateral thoracic contusion injury. Neither acute nor delayed treatment enhanced regeneration of CST axons into or past the lesion site. However, we observed significant sprouting of CST arbors in the cervical grey matter many segments rostral to the lesion after acute and chronic contusion in *Inpp5k* treated mice. These data cannot determine whether the sprouting axons were intact cervical CST axons or axotomized lumbar axons.

### Corticospinal neuron subtype specific therapies?

We profiled intact CSNs undergoing functional plasticity to identify novel intrinsic axon growth activators *Lppr1* and *Inpp5k*. Overexpression of both targets resulted in significant sprouting of CST axons into the cervical spinal cord, but not regeneration of axotomized CST axons after thoracic contusion injury. These data raise an important question regarding the homogeneity of CSNs. Our screen was completed in *crym*-GFP transgenic mice treated with retrograde tracers injected into the contralateral cervical enlargement. While the *crym*-GFP transgenic line is superior in labeling CSNs versus extrinsic tracers, it may not represent complete CSN coverage (Arlotta et al., 2005; Fink et al., 2015) and could preferentially label CSNs innervating the cervical cord. Thus, the bulk sequencing from our screen would have under sampled plastic lumbar CSNs. Additionally, we injected a retrograde tracer into the intact cervical spinal cord to identify CSNs whose arbors had sprouted across the midline. Thus, the combination of these approaches may have resulted in the identification of forelimb specific CSN growth activators. Indeed, we observed sprouting of CST axons in the cervical cord after uPyX, stroke and acute and chronic contusion injury. These data highlight the intriguing possibility that anatomically distinct CST terminal regions, such as the lumbar and cervical enlargements, may require disparate therapeutic interventions to stimulate axon growth. Therefore, a comprehensive understanding of the unique molecular profile of anatomical sub-regions of the CST is required to guide therapeutic interventions that require focus on specific regions of the spinal cord, for instance targeting lumbar CSNs after thoracic SCI.

In sum, we have demonstrated the potency of a second signaling pathway identified from our transcriptional screen of intact CSN undergoing functional plasticity to stimulate axon growth after acute and chronic CNS trauma. These data highlight the need to identify anatomically distinct subtypes of CSNs and determine whether these subtypes require unique therapeutic interventions to restore function after injury.

## Acknowledgements

This work was supported by grants from the CT Bioinnovations, The Craig Neilsen
Foundation, and the NIH.

